# Gain of gene regulatory network interconnectivity at the origin of vertebrates

**DOI:** 10.1101/2020.04.25.061077

**Authors:** Alejandro Gil-Gálvez, Sandra Jiménez-Gancedo, Rafael D. Acemel, Stephanie Bertrand, Michael Schubert, Héctor Escrivá, Juan J. Tena, José Luis Gómez-Skarmeta

**Affiliations:** Centro Andaluz de Biología del Desarrollo (CABD), Consejo Superior de Investigaciones Científicas-Universidad Pablo de Olavide-Junta de Andalucía, Seville, Spain; Sorbonne Université, CNRS, Biologie Intégrative des Organismes Marins, BIOM, Observatoire Océanologique, F-66650, Banyuls/Mer, France; Sorbonne Université, CNRS, Laboratoire de Biologie du Développement de Villefranche-sur-Mer, Institut de la Mer de Villefranche, Villefranche-sur-Mer, France

**Author notes:** These authors contributed equally (Alejandro Gil-Gálvez, Sandra Jiménez-Gancedo).

## Abstract

Signaling pathways control a large number of gene regulatory networks (GRNs) during animal development, acting as major tools for body plan formation^1^. Remarkably, in contrast to the large number of transcription factors present in animal genomes, only a few of these pathways operate during development^2^. Moreover, most of them are largely conserved along metazoan evolution^3^. How evolution has generated a vast diversity of animal morphologies with such a limited number of tools is still largely unknown. Here we show that gain of interconnectivity between signaling pathways, and the GRNs they control, may have played a critical contribution to the origin of vertebrates. We perturbed the retinoic acid, Wnt, FGF and Nodal signaling pathways during gastrulation in amphioxus and zebrafish and comparatively examined its effects in gene expression and cis-regulatory elements (CREs). We found that multiple developmental genes gain response to these pathways through novel CREs in the vertebrate lineage. Moreover, in contrast to amphioxus, many of these CREs are highly interconnected and respond to multiple pathways in zebrafish. Furthermore, we found that vertebrate-specific cell types are more enriched in highly interconnected genes than those tissues with more ancestral origin. Thus, the increase of CREs in vertebrates integrating inputs from different signaling pathways probably contributed to gene expression complexity and the formation of new cell types and morphological novelties in this lineage.

---

During embryonic development, thousands of genes are expressed in a coordinated and tightly regulated manner. This coordination is facilitated by complex hierarchical relationships between different genes^4,5^, where the expression of a determined gene triggers the transcription of many others, in a multi-level cascade that can involve hundreds of different genes^6^. Signaling pathways control most of these genetic cascades, interconnecting many genes and playing pivotal roles in more complex gene regulatory networks (GRNs). As a consequence, they are key substrates for the generation of morphological diversity during evolution^1^.

It has been already demonstrated that, after the vertebrate-specific whole genome duplications, many duplicated developmental genes were maintained in this group^7^. Furthermore, is also known that regulatory landscapes in general, and especially those of developmental genes, have been expanded in vertebrate linage^8^. However, it still remains unclear how these features interplay to generate organisms with higher complexity such as vertebrates. It is well known that the complexity of networks is not only dictated by the number of nodes but also by the number and patterns of interactions among those elements. In this context, effectors of signaling pathways constitute hubs in Gene regulatory networks (GRNs).

Here we investigate the contribution of key signaling pathways in the transition from invertebrates to vertebrates. To study this question, we compare the effect of interfering with the retinoic acid (RA), Wnt, FGF and Nodal pathways during gastrulation in both amphioxus (cephalochordate) and zebrafish embryos. To do so, we used compounds known to act either as agonists of the RA and Wnt or antagonists of the FGF and Nodal pathways^9^. We then examined the impact of these treatments on global gene expression by RNA-seq in both species (Fig. 1a, Extended Data Fig. 1). Additionally, we performed ATAC-seq to identify open chromatin regions^10^, including enhancers and promoters, affected by these manipulations (Fig. 1a, Extended Data Fig. 1).

**Figure 1.**
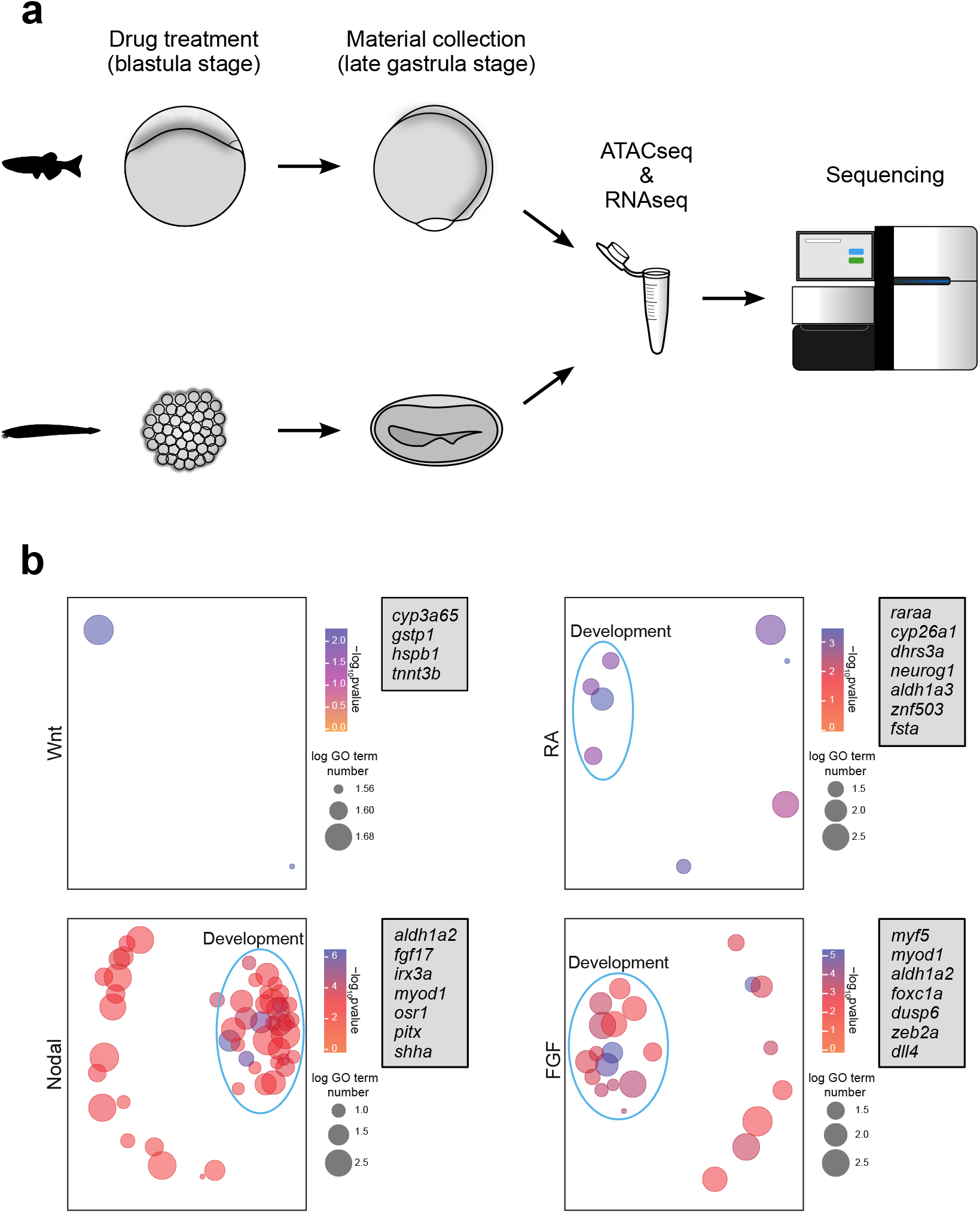
Experimental design and differential analyses. **a.** Overall design of the experiment: zebrafish and amphioxus embryos were treated with four different compounds at the blastula stage and dissociated at the late gastrula stage for RNA-seq and ATAC-seq library preparations. **b.** Gene ontology (GO) term enrichment analysis for common genes differentially expressed in both amphioxus and zebrafish. Each panel shows GO enrichment visualization obtained with the REVIGO tool for one of the treatments, taking into account only genes significantly modified upon the indicated treatment. Circles represent GO terms, and the X and Y axes map the semantic space: the closer the terms appear in the plot, the more related they are. In order to facilitate direct comparisons, a blue line surrounds GO terms associated with developmental processes. Gray boxes at the right of each panel show some of the developmental genes represented in the plot.

The whole genome duplications (WGDs) at the base of the vertebrate lineage^7,11^ and the additional WGD in the teleost lineage^12^ result in gene number imbalance between amphioxus and zebrafish. To overcome this limitation in our analysis, we used previously published data^8^ to retrieve all the vertebrate gene family members corresponding to each amphioxus gene affected by the treatments. For zebrafish, we only used the affected gene.

RNA-seq analysis revealed hundreds of differentially expressed genes following the different treatments in both amphioxus and zebrafish (Extended Data Fig. 1). Interestingly, we found transcripts similarly altered in both species upon the same treatment (Fig. 1b). Gene Ontology (GO) analysis for these common transcripts confirmed that they are highly associated with developmental processes, e.g. mesoderm, endoderm and hindbrain development, known to be regulated by the examined signaling pathways^9,13,14^ (Fig. 1b, Extended Data Table 1). Surprisingly, only few genes were similarly affected upon Wnt activation in both species (Fig. 1b, Extended Data Table 1). We then examined all the genes affected by each of these treatments, and we observed that the number of genes perturbed in zebrafish are higher than in amphioxus, and, proportionally, more strongly associated with development and signaling terms. Moreover, confirming our experimental approach, the genes altered by these treatments, and their corresponding GOs, clearly associate with developmental processes known to be regulated by these pathways^9,13,14^ (Extended Data Fig. 2, Extended Data Table 2).

We next analyzed our ATAC-seq data by searching for motifs enriched in peaks either more accessible after the treatment with RA or Wnt agonists, or less accessible upon the treatment with Nodal or FGF inhibitors. The binding sites for transcription factors (TFs) that mediate signaling by these pathways and/or well-known downstream TFs of these pathways were found for all treatments in both species^15–29^ (Extended Data Fig. 3, Extended Data Table 3). Overall, these results confirm that the pharmacological treatments of amphioxus and zebrafish embryos indeed perturbed the targeted pathways *in vivo*.

To better classify the genes that respond to interference of the different signaling pathways, we performed a clustering analysis of gene expression, which resulted in groups of genes with similar transcriptional behavior (Fig. 2a). We then carried out GO enrichment analyses of these groups. The RNA-seq-derived clusters in zebrafish were mostly associated with GO terms related to embryonic development, while in amphioxus we also detected many terms related to metabolism and cell homeostasis (Fig. 2a, Extended Data Table 4). In some cases, the GO terms were specifically associated with a single pathway. In amphioxus, this was for example the case for retinol metabolism in genes upregulated by RA treatment (dark blue cluster) and for muscle cell differentiation in genes downregulated by Nodal inhibition (orange cluster). In zebrafish, heart formation and somitogenesis were associated with, respectively, Nodal and FGF inhibition (orange and light blue clusters), while brain development was associated with Wnt pathway activation (light green cluster) (Fig. 2a). This cluster analysis also revealed that, in both amphioxus and zebrafish, the majority of developmental processes are influenced by several of the studied signaling pathways.

**Figure 2.**
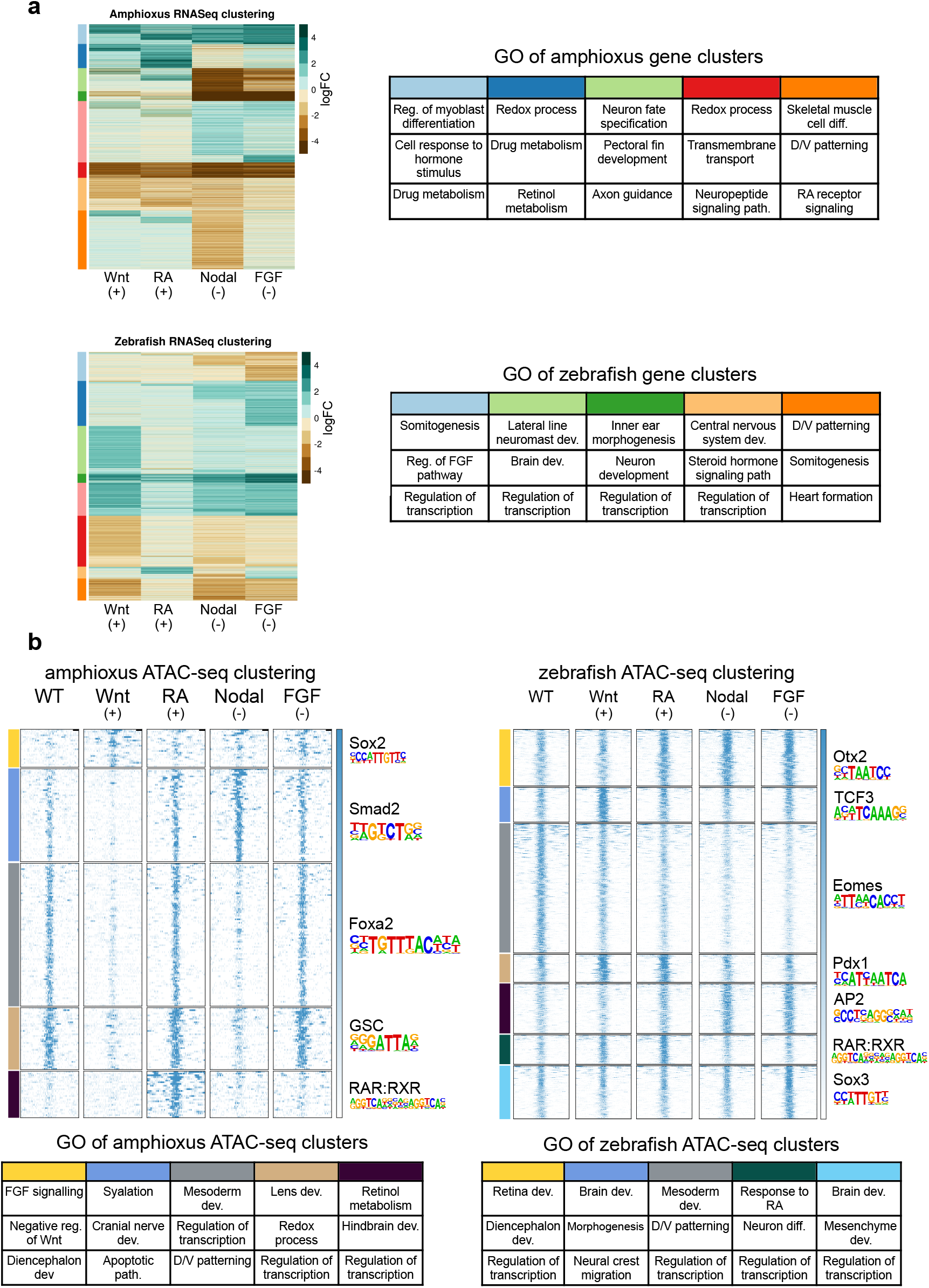
Clustering of RNA-seq and ATAC-seq data. **a.** RNA-seq-derived gene expression fold change (FC) in amphioxus (upper panel) and zebrafish (bottom panel). FC values are calculated for each condition versus control samples by means of DGE analyses. Tables on the right show three representative gene ontology (GO) terms enriched in the cluster marked with the same color. **b.** Clustering of ATAC-seq data in amphioxus (left) and zebrafish (right). Top DNA binding motif found in each cluster is represented on the right, and three representative GO terms enriched in the clusters are shown in the tables below.

In the case of ATAC-seq peaks clustering, we assigned peaks to their putative target genes using the GREAT algorithm^30^, in order to derive GO terms. In general, we observed a good correlation between the average ATAC-seq signal around the peaks in the different clusters, the GO terms of their putative target genes and the TF binding motifs identified within these peaks (Fig. 2b). Furthermore, open chromatin regions altered upon treatments were mainly associated with development, especially in zebrafish (Fig. 2b), which is consistent with our RNA-seq analysis. For example, the zebrafish cluster of ATAC-seq peaks that was, in average, more accessible upon Wnt stimulation (blue cluster), was associated with brain development (Fig. 2b). In addition, TCF3 motifs were found within these peaks (Fig. 2b). Similarly, in both species, the clusters enriched following RA treatment (dark purple cluster in amphioxus, dark green cluster in zebrafish) were associated with response to RA and hindbrain development (Fig. 2b). The clusters in both species further contain RAR:RXR motifs (Fig. 2b).

The majority of ATAC-seq peaks clusters are characterized by changes in more than one signaling pathway, which is similar to what we observed for the RNA-seq clustering. Several signaling pathways thus seem to act on the same ATAC-seq peaks, suggesting a certain level of interconnection between the different pathways.

In order to directly compare the integrated effect of the treatments at the transcriptomic and regulatory levels in both species, we intersected the differentially regulated genes at the transcriptomic level with the genes associated with differential ATAC-seq peaks for each treatment and species. In this analysis, we included both positively and negatively affected genes/regulatory elements. The intersection resulted in a total of 2098 genes and 4609 ATAC-seq peaks in zebrafish and 481 genes and 853 ATAC-seq peaks in amphioxus (Extended Data Table 5). Although the lower numbers of treatment-affected genes and ATAC-seq peaks in amphioxus could correspond to a loss of regulatory information in this species, it is known that there was a general gain of regulatory input in vertebrates^8^. Thus, it is very likely that this rather indicates a gain of response to these signaling pathways in vertebrates, through the incorporation of novel cis-regulatory elements (CREs).

We then clustered the results of GO enrichment analyses associated with these genes in amphioxus and zebrafish (Fig. 3, Extended Data Fig. 4). We found that the p-values associated to the enriched GO terms were, in general, much lower in zebrafish than in amphioxus, and the number of GO terms significantly affected by the different treatments were much higher (Fig. 3). This suggests that, in the vertebrate, the regulatory networks involved in developmental processes are more complex. The same effect was observed using the number of gene families (which are not affected by the overestimation of the number of genes in amphioxus due to the inclusion of whole families of orthologous genes in our lists) instead of p-values (Extended Data Fig. 4).

**Figure 3.**
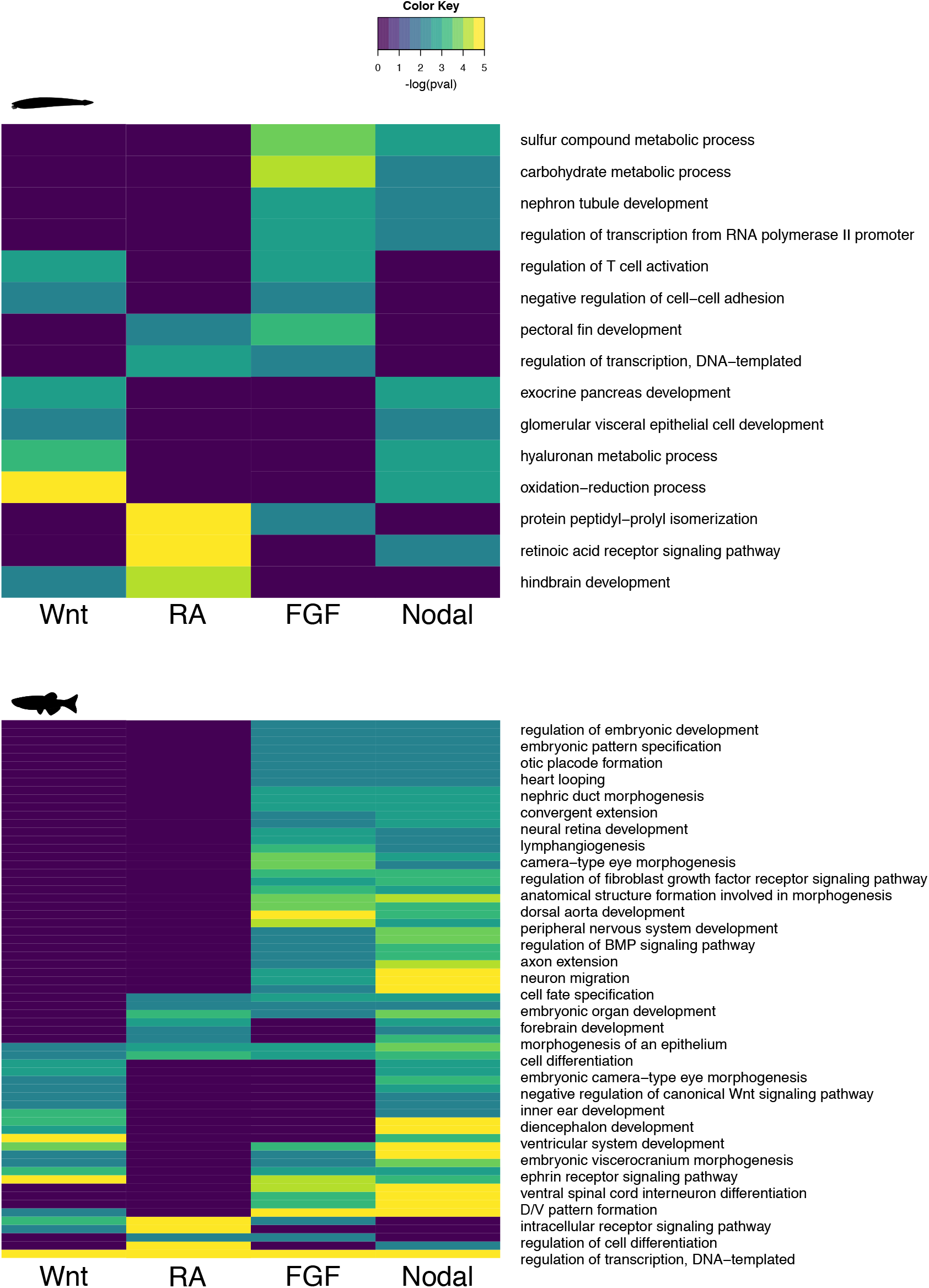
Clustering of GO terms. Gene ontology (GO) terms enriched in differentially regulated genes at the transcriptomic level (based on RNA-seq analysis) that are also associated to differential ATAC-seq peaks, clustered by associated p-values (pval) in amphioxus (upper panel) and zebrafish (bottom panel).

These results further support that there are more genes controlled by these signaling pathways in vertebrates. Among the genes we identified in zebrafish were a large number of TFs and regulators of different signaling pathways (Extended Data Table 6). We also found that some GO terms in zebrafish were significantly enriched for signaling pathways (for example, FGF and Nodal pathways share many development-related terms, Fig. 3). These two facts indicate that, in agreement with the results of our previous clustering analysis, different signaling pathways are interconnected and that the degree of connectivity is higher in vertebrates compared to invertebrates. To directly test this, we counted the number of genes that were affected by manipulating one, two, three or four different signaling pathways in zebrafish and amphioxus. Interestingly, independently of considering only RNA-seq data or combining RNA-seq and ATAC-seq information, we found that the interconnection between pathways was higher in zebrafish than in amphioxus, with a higher proportion of genes responding to two, three or four perturbations in zebrafish (Fig. 4a, Extended Data Fig. 5a). Moreover, we observed an increase of connectivity in vertebrates independent of the number of genes retained after the WGD events (Extended Data Fig. 5b). Nevertheless, the more copies retained, the higher the gain of connectivity (Extended Data Fig. 5c). Using already available single cell RNA-seq data^31^ we observed that, interestingly, tissues that appeared as novelties during the invertebrate to vertebrate transition, such as the neural crest cells or the sensory placodes, showed an enrichment in the expression of highly connected genes, while in more evolutionary ancient tissues, like the muscles or the intestine, the expression of highly and lowly connected genes was very similar (Fig. 4b, Extended Data Fig. 6). Finally, we used the Cytoscape tool^32^ to visualize all connections between genes and the different signaling pathways in amphioxus and zebrafish (Fig. 4c). This plot clearly shows that developmental GRNs associated with these four signaling pathways are more interconnected in vertebrates.

**Figure 4.**
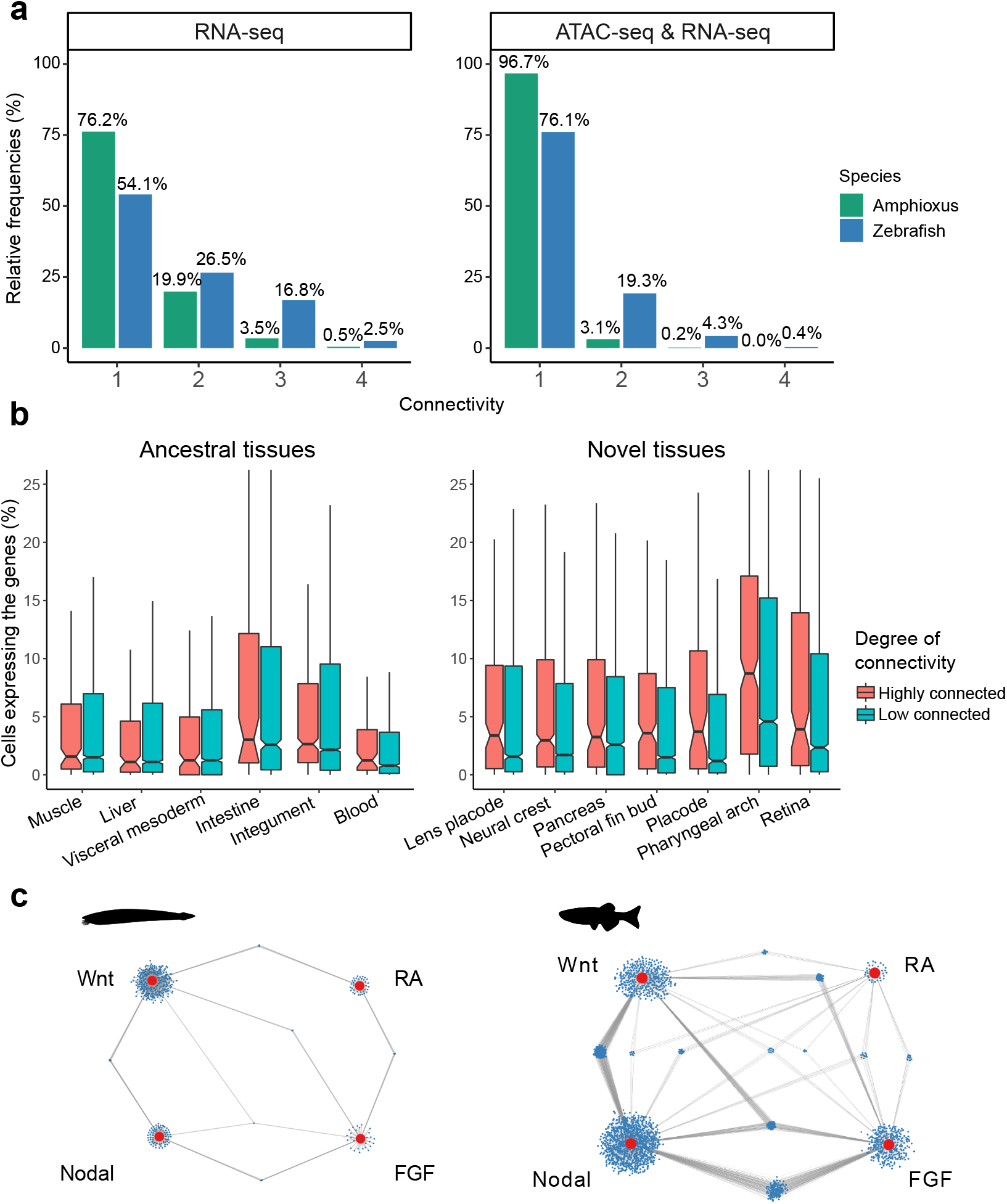
Gene connectivity to different signaling pathways. **a.** Percentage of genes that respond to one, two, three or four different pathways in amphioxus (green bars) and zebrafish (blue bars), only at the RNA-seq level (left panel) or at both the RNA-seq and ATAC-seq level (right panel). **b.** Number of cells expressing highly connected genes (connectivity >= 3, pink bars) and lowly connected genes (connectivity = 1, light blue bars) according to single cell RNA-seq experiments carried out in zebrafish^31^ in ancestral tissues (and thus also present in amphioxus) (left panel) and in vertebrate specific tissues (right panel). **c.** Cytoscape plot showing connectivity networks in amphioxus (left panel) and zebrafish (right panel). Small blue dots represent genes and big red dots represent signaling pathways. Gray lines mark the responsiveness of each gene to the connected signaling pathway.

By comparing epigenomic and transcriptomic data in amphioxus, zebrafish and other vertebrates, we recently showed that the invertebrate to vertebrate transition is associated with an increase of regulatory information in the latter lineage and that this increase likely contributed to the spatial and functional specialization of some duplicated genes^8^. Here, we demonstrate that some of the novel vertebrate CREs contribute to a more complex interconnection between the RA, Wnt, FGF and Nodal signaling pathways. An increased interconnection between these four key developmental signaling pathways likely facilitated the restriction of the expression domains of some duplicated developmental genes, which, in turn, contributed to the increment of tissue complexity required to generate morphological novelties in vertebrates.

## Acknowledgements

We thank all members of JLGS laboratory for fruitful discussions, and C3UPO for HPC support. We especially thank José María Santos-Pereira, Martin Franke, Christina Paliou Ignacio Maeso and Manuel Irimia for helpful comments on the manuscript. This project has received funding from the European Research Council (ERC) under the European Union’s Horizon 2020 research and innovation programme (grant agreement No 740041), the Spanish Ministerio de Economía y Competitividad (grants BFU2016-74961-P to JLGS and BFU2014-58449-JIN to JJT). This work was also supported by the institutional grant Unidad de Excelencia María de Maeztu (MDM-2016-0687 to the Department of Gene regulation and morphogenesis of Centro Andaluz de Biología del Desarrollo). HE and SB were supported by the CNRS and the ANR CHORELAND, ANR-16-CE12-0008-01.

## Author contributions

A.G.-G., S.J.-G., J.J.T. and R.D.A. performed experiments and computation analyses. S.B. and H.E. obtained biological material and performed experiments. M.S. provided reagents. J.J.T. and J.L.G.-S. coordinated the project, contributed to the study design and wrote the main text with input from all authors.

## Methods

### Animal husbandry and treatment of embryos

*Danio rerio* embryos were manipulated following the protocols that have been approved by the Ethics Committee of the Andalusia Government (license number 182-41106) and the national and European regulation established. All experiments with zebrafish were carried out in accordance with the principles of 3Rs (replacement, reduction and refinement). Zebrafish embryos at 30% epiboly were treated by adding the different drugs to the medium. In the case of SB505124, embryos were treated at 256 cells stage. The final concentrations of drugs were: ATRA 0.1μM (cat n° R2625, Sigma-Aldrich Merck), SU5402 20μM (cat n° SML0444, Sigma-Aldrich Merck), BIO 14μM (cat n° B1686, Sigma-Aldrich Merck) and SB505124 30μM (cat n° S4696, Sigma-Aldrich Merck). These molecules were dissolved in DMSO (cat n° D2438, Sigma-Aldrich Merck). As a control experiment, the same volume of DMSO was added to the medium. Drugs were removed by changing the medium several times when embryos were at 80% epiboly stage. After that, embryos were carefully transferred to a glass Petri dish and chorion was removed using pronase.

*Branchiostoma lanceolatum* adults were collected at the Racou beach in Argelès-sur-Mer (France). Spawning was induced as previously described^33,34^ and the fertilization of eggs was done in vitro. Amphioxus embryos were manipulated in filtered seawater unless otherwise specified. After fertilization, embryos were dechorionated in a Petri dish covered with agarose (0.8% agarose in filtered seawater) by pippeting them towards the border of the dish, and gently transferred to a small Petri dish. Drugs were dissolved in DMSO at the following final concentrations: ATRA 1μM (cat n° R2625, Sigma-Aldrich Merck), SU5402 25μM (cat n° 572630, Sigma-Aldrich Merck), BIO 1μM (cat n° B1686, Sigma-Aldrich Merck) and SB505124 50μM (cat n° S4696, Sigma-Aldrich Merck). SB505124 drug was added to the filtered seawater at 3 hours post-fertilization stage, while the rest of the drugs were added at 5 hours post-fertilization stage. Control embryos were treated with 0.1% DMSO. Finally, at 15 hours post-fertilization, treated embryos were washed several times with filtered seawater in order to remove the drugs from the medium.

### ATAC-seq

ATAC-seq assays were performed at least in two biological replicates. 45 Amphioxus embryos and 20 zebrafish embryos were dissociated in individual cells. After counting the number of cells, around 70000 cells were transferred to another tube to perform the experiment. ATAC-seq experiments were performed as previously described^8,10^.

ATAC-seq data analyses were performed using standard pipelines^8,10^. Reads were aligned with Bowtie2 using GRCz10 (danRer10) and Bl71^8^ assemblies for zebrafish and amphioxus samples, respectively. Those reads that were separated by more than 2 Kb were filtered out of the analysis. The exact position of the Tn5 cutting site was determined as the position −4 (reverse strand) or +5 (forward strand) from the read start, and this position was extended 5 Kb in both directions. BED files were transformed into BigWig using the wigToBigWig UCSC tool. Reads were extended 100 bp in order to visualize the data in the UCSC Genome Browser^35^. Macs2^36^ software was used in order to perform the peak calling in each sample, using the parameters–nomodel, --shift 45 and --extsize 100. The different called peaks of each sample were merged into a unique set of peaks, taking into account replicates (two per sample). Then, using Bedtools^37^, we computed the number of reads per called peak and per sample in both treatment and control conditions, and a differential analysis was performed using DESeq2^38^ v1.18.0 in R 3.4.3. A corrected p value < 0.05 was set as cutoff for statistical significance of the differential accessibility of the chromatin in ATAC-seq peaks. Motif enrichment was calculated using the program FindMotifsGenome.pl from Homer tool suite^39^.

k-means clustering of ATAC-seq signal was performed using Deeptols 2.0^40^ and seqMiner^41^.

The assignment of ATAC-seq peaks to genes was done using the GREAT tool^42^, with default parameters for basal plus extension regions calculation. The Gene Ontology analysis was carried out using TopGO R package, using the *elim* test for taking the most specific GO enriched terms.

### RNA-seq

RNAseq experiments were performed in three biological replicates for each species. RNA samples were extracted from 15 zebrafish and 100 amphioxus embryos following already published protocols^8^.

For the data analysis, reads were aligned against GRCz10 (danRer10) and Bl71 assemblies using STAR^43^ v2.5.3a and assigned to genes using the HTSeq toolkit^44^ v0.11.2. Differential gene expression analyses were performed using DESeq2^38^ v1.18.0. A corrected p value < 0.05 and an absolute log_2_FC > 1 were used as thresholds for calling differential genes. The enrichment of Biological process GO terms was calculated using the TopGO R package.

Gene clusterings were performed using Pheatmap 1.0.12 R package, using kmeans as clustering method.

The integration of differential ATAC-seq peaks and differential expressed genes was done at gene level in both species. The gene lists that resulted from both analyses for the same treatment were intersected in order to find genes that had a differential RNA-seq signal and also one or more associated differential ATAC-seq peaks.

Since there is no functional annotation of the amphioxus genes, the orthologous zebrafish genes were used for computing the GO term enrichment analysis. Amphioxus *vs*. zebrafish orthologous genes were already available from previous studies^8^ (Extended Data Table 7).

### Connectivity analyses

In order to categorize genes that could be responding to more than one treatment, we computed a connectivity score. For this, we took into account only genes that responded at both ATAC-seq and RNA-seq levels. Each gene was assigned with a discrete score that ranges from 1 to 4, corresponding to the number of different treatments that they responded to. Cytoscape^32^ networks were generated in order to better represent the connectivity of responsive genes. Connectivity tables are available in Extended Data Table 7.

### scRNA-seq analysis

We selected genes with high connectivity (>= 3) and low connectivity (=1) from the previously published scRNA-seq zebrafish developmental atlas^31^ and explore their expression levels in a set of ancient and vertebrate-specific novel tissues. A gene is defined as expressed in a cell if its normalized expression in that cell is greater than 0. For a certain tissue, we calculated the proportion of cells that expressed a certain gene, and aggregated these proportions for those genes that have high or low connectivity score in a boxplot. In order to determine which genes were expressed in each tissue, we set a threshold of a 5% of the cells of that specific tissue. Subsequently, for a direct comparison between a set of ancestral tissues, present in both zebrafish and amphioxus, and another group of vertebrate-specific tissues, we computed and represented in barplots the ratio between genes with high connectivity score and those with low score expressed in each tissue.

## References

1. Pires-daSilva, A. & Sommer, R. J. The evolution of signalling pathways in animal development. Nature Reviews Genetics 4, 39–49 (2003).

2. Sanz-Ezquerro, J. J., Münsterberg, A. E. & Stricker, S. Editorial: Signaling pathways in embryonic development. Frontiers in Cell and Developmental Biology 5, 76 (2017).

3. Babonis, L. S. & Martindale, M. Q. Phylogenetic evidence for the modular evolution of metazoan signalling pathways. Philosophical Transactions of the Royal Society B: Biological Sciences 372, (2017).

4. Ravasz, E., Somera, A. L., Mongru, D. A., Oltvai, Z. N. & Barabási, A. L. Hierarchical organization of modularity in metabolic networks. Science (80-.). 297, 1551–1555 (2002).

5. Barabási, A. L. & Oltvai, Z. N. Network biology: Understanding the cell’s functional organization. Nature Reviews Genetics 5, 101–113 (2004).

6. Azpeitia, E. et al. The combination of the functionalities of feedback circuits is determinant for the attractors’ number and size in pathway-like Boolean networks. Sci. Rep. 7, 42023 (2017).

7. Putnam, N. H. et al. The amphioxus genome and the evolution of the chordate karyotype. Nature 453, 1064–1071 (2008).

8. Marlétaz, F. et al. Amphioxus functional genomics and the origins of vertebrate gene regulation. Nature 564, 64–70 (2018).

9. Bertrand, S., Petillon, Y. Le, Somorjai, I. M. L. & Escriva, H. Developmental cell-cell communication pathways in the cephalochordate amphioxus: Actors and functions. Int. J. Dev. Biol. 61, 697–722 (2017).

10. Buenrostro, J. D., Giresi, P. G., Zaba, L. C., Chang, H. Y. & Greenleaf, W. J. Transposition of native chromatin for fast and sensitive epigenomic profiling of open chromatin, DNA-binding proteins and nucleosome position. Nat. Methods 10, 1213–1218 (2013).

11. Dehal, P. & Boore, J. L. Two rounds of whole genome duplication in the ancestral vertebrate. PLoS Biol. 3, e314 (2005).

12. Christoffels, A. et al. Fugu genome analysis provides evidence for a whole-genome duplication early during the evolution of ray-finned fishes. Mol. Biol. Evol. 21, 1146–1151 (2004).

13. Kiecker, C., Bates, T. & Bell, E. Molecular specification of germ layers in vertebrate embryos. Cellular and Molecular Life Sciences 73, 923–947 (2016).

14. Tuazon, F. B. & Mullins, M. C. Temporally coordinated signals progressively pattern the anteroposterior and dorsoventral body axes. Semin. Cell Dev. Biol. 42, 118–133 (2015).

15. Böttcher, R. T. & Niehrs, C. Fibroblast growth factor signaling during early vertebrate development. Endocr. Rev. 26, 63–77 (2005).

16. Cadigan, K. M. & Waterman, M. L. TCF/LEFs and Wnt signaling in the nucleus. Cold Spring Harb. Perspect. Biol. 4, (2012).

17. Charney, R. M., Paraiso, K. D., Blitz, I. L. & Cho, K. W. Y. A gene regulatory program controlling early Xenopus mesendoderm formation: Network conservation and motifs. Seminars in Cell and Developmental Biology 66, 12–24 (2017).

18. Bian, S. S. et al. Clock1a affects mesoderm development and primitive hematopoiesis by regulating Nodal-Smad3 signaling in the zebrafish embryo. J. Biol. Chem. 292, 14165–14175 (2017).

19. Jia, S., Ren, Z., Li, X., Zheng, Y. & Meng, A. smad2 and smad3 are required for mesendoderm induction by transforming growth factor-ß/nodal signals in zebrafish. J. Biol. Chem. 283, 2418–2426 (2008).

20. Kjolby, R. A. S., Truchado-Garcia, M., Iruvanti, S. & Harland, R. M. Integration of Wnt and FGF signaling in the xenopus gastrula at TCF and Ets binding sites shows the importance of short-range repression by TCF in patterning the marginal zone. Dev. 146, (2019).

21. Tian, T. & Meng, A. M. Nodal signals pattern vertebrate embryos. Cell. Mol. Life Sci. 63, 672–685 (2006).

22. Friedman, J. R. & Kaestner, K. H. The Foxa family of transcription factors in development and metabolism. Cell. Mol. Life Sci. 63, 2317–2328 (2006).

23. Ghyselinck, N. B. & Duester, G. Retinoic acid signaling pathways. Dev. 146, (2019).

24. Hoodless, P. A. et al. FoxH1 (Fast) functions to specify the anterior primitive streak in the mouse. Genes Dev. 15, 1257–1271 (2001).

25. Joshi, P., Darr, A. J. & Skromne, I. CDX4 regulates the progression of neural maturation in the spinal cord. Dev. Biol. 449, 132–142 (2019).

26. Kjolby, R. A. S. & Harland, R. M. Genome-wide identification of Wnt/ß-catenin transcriptional targets during Xenopus gastrulation. Dev. Biol. 426, 165–175 (2017).

27. Aldea, D. et al. Genetic regulation of amphioxus somitogenesis informs the evolution of the vertebrate head mesoderm. Nat. Ecol. Evol. 3, 1233–1240 (2019).

28. Onai, T. Canonical Wnt/ß-catenin and Notch signaling regulate animal/vegetal axial patterning in the cephalochordate amphioxus. Evol. Dev. 21, 31–43 (2019).

29. Yasuoka, Y., Tando, Y., Kubokawa, K. & Taira, M. Evolution of cis-regulatory modules for the head organizer gene goosecoid in chordates: Comparisons between Branchiostoma and Xenopus. Zool. Lett. 5, 27 (2019).

30. McLean, C. Y. et al. GREAT improves functional interpretation of cis-regulatory regions. Nat. Biotechnol. 28, 495–501 (2010).

31. Farnsworth, D. R., Saunders, L. M. & Miller, A. C. A single-cell transcriptome atlas for zebrafish development. Dev. Biol. 459, 100–108 (2020).

32. Shannon, P. et al. Cytoscape: A software Environment for integrated models of biomolecular interaction networks. Genome Res. 13, 2498–2504 (2003).

## References for Methods section

33. Fuentes, M. et al. Preliminary observations on the spawning conditions of the European amphioxus (Branchiostoma lanceolatum) in captivity. J. Exp. Zool. Part B Mol. Dev. Evol. 302, 384–391 (2004).

34. Fuentes, M. et al. Insights into spawning behavior and development of the European amphioxus (Branchiostoma lanceolatum). J. Exp. Zool. Part B Mol. Dev. Evol. 308, 484–493 (2007).

35. Casper, J. et al. The UCSC Genome Browser database: 2018 update. Nucleic Acids Res. 46, D762–D769 (2018).

36. Zhang, Y. et al. Model-based analysis of ChIP-Seq (MACS). Genome Biol. 9, R137 (2008).

37. Quinlan, A. R. & Hall, I. M. BEDTools: A flexible suite of utilities for comparing genomic features. Bioinformatics 26, 841–842 (2010).

38. Love, M. I., Huber, W. & Anders, S. Moderated estimation of fold change and dispersion for RNA-seq data with DESeq2. Genome Biol. 15, (2014).

39. Heinz, S. et al. Simple Combinations of Lineage-Determining Transcription Factors Prime cis-Regulatory Elements Required for Macrophage and B Cell Identities. Mol. Cell 38, 576–589 (2010).

40. Ramírez, F. et al. deepTools2: a next generation web server for deep-sequencing data analysis. Nucleic Acids Res. 44, W160–W165 (2016).

41. Ye, T. et al. seqMINER: An integrated ChIP-seq data interpretation platform. Nucleic Acids Res. 39, e35–e35 (2011).

42. Hiller, M. et al. Computational methods to detect conserved non-genic elements in phylogenetically isolated genomes: Application to zebrafish. Nucleic Acids Res. 41, (2013).

43. Dobin, A. et al. STAR: Ultrafast universal RNA-seq aligner. Bioinformatics 29, 15–21 (2013).

44. Anders, S., Pyl, P. T. & Huber, W. HTSeq-A Python framework to work with high-throughput sequencing data. Bioinformatics 31, 166–169 (2015).

